# Plasticity of the gastrocnemius elastic system in response to decreased work and power demand during growth

**DOI:** 10.1101/2021.04.12.439563

**Authors:** SM Cox, A DeBoef, MQ Salzano, K Katugam, SJ Piazza, J Rubenson

## Abstract

Elastic energy storage and release can enhance performance that would otherwise be limited by the force-velocity constraints of muscle. While functional influence of a biological spring depends on tuning between components of an elastic system (the muscle, spring, driven mass, and lever system), we do not know whether elastic systems systematically adapt to functional demand. To test whether altering work and power generation during maturation alters the morphology of an elastic system, we prevented growing guinea fowl (*Numida Meleagris*) from jumping. At maturity, we compared the jump performance of our treatment group to that of controls and measured the morphology of the gastrocnemius elastic system. We found that restricted birds jumped with lower jump power and work, yet there were no significant between-group differences in the components of the elastic system. Further, subject-specific models revealed no difference in energy storage capacity between groups, though energy storage was most sensitive to variations in muscle properties (most significantly operating length and least dependent on tendon stiffness). We conclude that the gastrocnemius elastic system in the guinea fowl displays little to no plastic response to decreased demand during growth and hypothesize that neural plasticity may explain performance variation.

## Introduction

Taking advantage of storage and release of elastic strain energy can enhance performance that would otherwise be limited by the force-velocity constraints of muscle. The temporal decoupling of energy production from energy delivery permitted by elastic energy storage allows muscles and tendons to produce force effectively over a range of shortening or lengthening speeds. Muscles may generate forces during slow or isometric contractions and elastic recoil augments the rate of energy delivery or absorption during rapid movements [1,2]. By making use of energy storage in the tendon “spring”, a muscle tendon unit (MTU) can produce force more economically or with greater power than a muscle alone [2]. Yet, several studies have identified important differences among spring-muscle combinations. Wilson, Lichwark, and colleagues [3,4] showed that the efficiency of an MTU during cyclic loading depends on the tuning of relative muscle and spring properties. For instance, muscle efficiency varies with both fascicle length and tendon stiffness, with the specific optimal efficiency values depending on gait conditions [3]. Several researchers [5–9] have shown that the opposing inertial or drag forces acting on a motor-spring system also influence whether springs enhance performance. Adding further complexity, Sawicki and colleagues [10,11] found that the timing of neural activation of muscle during hopping must be tightly controlled to take advantage of in-series springs. Together, this body of work suggests that understanding the conditions in which spring systems enhance performance may require expanding our focus from the muscle-tendon unit to that of the broader ‘elastic system’ which includes the muscle (motor), the spring, the resistive forces, and the neural control of the system. The optimal performance of an elastic system may require tuning of both morphology and neural control. This approach recognizes the integrated nature of the neuro-musculoskeletal system[12].

The sensitivity of elastic system efficiency to the tuning of its components complicates inferences for how elastic systems systematically adapt to functional demand during maturation. For instance, do growing individuals who regularly perform functions that utilize elastic strain energy develop elastic systems with greater energy storage capacity? This is still unknown because most studies of MTU plasticity have focused on how individual components of elastic systems (neural control [13– 15], muscle [16–19] and tendon [20–25]) vary with task or training, and how those individual changes influence function of a muscle-tendon unit [10,11,26–28]. Yet, the integrated nature of the elastic system suggests that functional consequences of plasticity are difficult to predict by analyzing elements in isolation [27,29,30]. Therefore, the complex nature of the neuromuscular adaptation of elastic systems may require analysis at the system level rather than at the level of individual components.

Here we present a study of the morphological plasticity of an elastic system. Specifically, we ask whether individuals that jump during maturation (an activity requiring elastic energy storage and return [31]) develop elastic systems that more capable of storing elastic strain energy at maturity than those of individuals restricted from jumping. Here we focus on the elastic system most involved in storage and release of elastic energy during jumping[32–36], the gastrocnemius elastic system. We test this by altering the rearing conditions of two groups of guinea fowl (*Numida meleagris*) across the entire growth period, allowing one group to engage in jump-to-perch behavior and preventing all jumping in the other group. We previously reported that restricted birds in this study showed detriments in jump performance at adulthood [37]. In this manuscript, we aim to link the morphological and functional consequences of our intervention.

Of the morphology data, we take both an individual-component and systems-level approach to evaluate the plasticity of an elastic system during growth. At the component level, we probe whether our treatment resulted in systematic morphological differences in individual components of the gastrocnemius elastic system between groups. We seek to determine whether components of this elastic system plastically adapt to variations in functional demand during growth. At the systems level, we ask how plastic changes at the component level interact to influence the capacity for elastic energy storage. To do this, we developed subject-specific musculoskeletal models that incorporated experimentally measured morphological properties of each bird’s elastic system. With each subject-specific model, we simulated a fully activated muscle contraction under various postures and quantified the resulting tendon energy stored. The purpose of this systems-level analysis was to evaluate the integrated effects of morphological variation.

The component-and systems-level analyses serve as a case study for understanding how a particular elastic system changes with functional demand. We also took advantage of the variation within and across groups to ask broader questions about the relationship between form and function in elastic systems. Specifically, we asked which combinations of naturally occurring morphological variation most influence the ability of an elastic system to store energy. Lastly, we probed the extent to which energy storage capacity in the gastrocnemius elastic system correlated with jump performance.

We hypothesize that components of an elastic system plastically adapt to variations in functional demand during maturation, resulting in greater energy storage capacity in birds that jump during growth. We predict energy storage capacity will increase linearly with muscle force-generating capacity and inversely with tendon stiffness [38–40]. Finally, we predict that differences in jump performance positively correlate with an animal’s ability to store elastic strain energy in the tendon of the gastrocnemius elastic system.

## Methods

### Experimental Protocol

#### Animals

To study these questions, one-day-old guinea fowl keets (*Numida meleagris*) were obtained from a regional breeder (Guinea Farm; New Vienna, IA). After a 2-wk brooding period, the keets were pen reared through skeletal maturity (>6 months) in one of two conditions, as we previously described in detail (Cox et al 2020). A control group (C; n =8) was housed in a large, circular pen (3.14 m^2^) that allowed ample room for locomotion and objects for jumping and perching. The restricted treatment group (R; n = 7) were raised a smaller pen (1 m^2^ at maturity) with low mesh ceilings that prevented jumping. Food and water were available *ad libitum* (food intake did not differ between groups). Lights were programmed to be on a 12:12-h light-dark cycle. The experimental protocol was approved by Institutional Animal Care and Use Committee at The Pennsylvania State University (IACUC; Ref. #46435).

#### Movement Analysis

As described previously [37], to quantify the influence of pen configuration on the movement patterns of both treatment groups, pens were filmed from above for ten minutes, four times per day, across the growth period (Foscam; C2 1080p HD cameras, Houston, TX). For each bird in the pen, behaviors were tracked and categorized into two states (standing or walking) and three events [short sprint (<2 sec), hurdle jump (< body height) and perch jump (∼2 x body height).

### Functional Measures

As described previously [37], jump performance was measured by placing each bird in turn on 6×6 in. force plates (AMTI HE6×6; Watertown, MA, USA) enclosed in a tapered box and encouraging the birds to jump. Jump power was calculated from the instantaneous net vertical ground reaction and the vertical center of mass velocity. Velocity was obtained by integrating the center of mass acceleration, which was in turn found from the net ground reaction force and the body mass. We calculated jump work by integrating the instantaneous power with respect to time over the course of the jump.

### Quantification of properties of individual components of the elastic system

#### Specimen muscle architecture preparation

The pelvic limb was separated from the upper body and the left and right legs were then split by sectioning the pelvis at the midline while avoiding muscle attachments. Right limbs were placed into neutral buffered formalin for fixation (10%) for at least two weeks, while left legs were fresh-frozen and kept at −20 °C. Right limbs were positioned with joint angles approximating those at mid-swing during running (hip: 30°, knee: 80°, ankle: 125°, within ±2° [41]). Joint angles were confirmed for the fixed limbs using photographs made with a digital camera (Canon EOS550D; Surrey, United Kingdom) and analyzed with ImageJ (National Institutes of Health, Betesda, MD). Left limbs were fresh-frozen and saved for later muscle mass measures.

#### Muscle Analyses

We made measurements of the lateral and medial heads of the gastrocnemius muscle (LG and MG), the muscle group of the MTU thought primarily responsible for storage and release of elastic strain energy during running and jumping [31,36]. The third (intermedia) head of the gastrocnemius only comprises ∼10% of the total mass of the gastrocnemius muscles in this species [42] and thus was not included in the analysis. MG and LG were dissected from the fresh-frozen left limbs and weighed to the nearest 0.1 milligram. The LG and MG were then dissected from the fixed limbs for fascicle length, pennation angle, and sarcomere analysis. LG was first split longitudinally through the mid-belly to view fascicle arrangement. Photographs of whole MG and split LG made with a digital camera (Canon EOS550D) were imported into ImageJ for measurement of the pennation angle between muscle fascicles and their insertions on the aponeurosis [43].

Due to the expected within-muscle heterogeneity of strain [44,45], each muscle was divided into sections for analysis. MG was split into anterior and posterior fascicles [46] and then again split proximally/distally, resulting in four sections. The LG was split into proximal, middle, and distal sections, each spanning one-third the length of the muscle belly. Average pennation angle was found for each section by taking the mean of three angle measurements. Sarcomere lengths for each section were found using the laser diffraction techniques described in [43]. A minimum of three sarcomere length measurements were taken from each muscle fascicle bundle and these measurements were averaged to obtain the mean measured sarcomere length.

Optimal fascicle length, L_O_, was calculated by multiplying the length of the fascicle by the ratio of optimal sarcomere length of guinea fowl muscle (2.36 μm; [46]) to the mean measured sarcomere length.

Pennation angle at optimal fascicle length, *θ*_*OFL*_, was calculated from the average measured pennation angle, 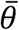, and the ratio of measure fiber length, *F*_*l*_, and calculated optimal fascicle length, L_O_ according to the equation [47]: 

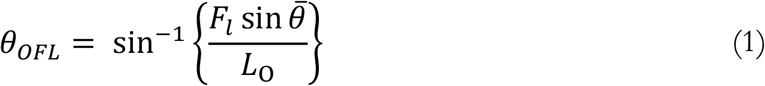

Maximum isometric force along the muscle fiber for the MG and LG were calculated from the muscle mass, *m*, optimal fascicle length, *L*_*O*_ and muscle density (ϱ_musc_ = 1060 kg/m^3^ [48] using the specific tension, *f* (3 x 10^5^ N/m^2^, Rospars and Meyer-Vernet, 2016), according to the equation: 

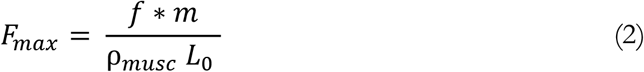

We specifically calculated isometric force along the muscle fiber for input into the musculoskeletal model rather than including the influence of pennation angle because pennation angle is a separate input into the OpenSim Millard muscle model (see below for model description), which accounts for the change in pennation angle with muscle length [50]. For statistical tests, muscle force along the tendon was analyzed.

#### Moment Arm

Gastrocnemius moment arm at the ankle was experimentally measured using the tendon travel method [51,52] as described by Salzano (2020). The gastrocnemius moment arm at the ankle was experimentally measured using the tendon travel method as described by Salzano [53]. In short, the Achilles tendon was attached to a linear transducer (Model P510-2-S11-N0S-10C, UniMeasure, Inc., Corvallis, OR) to measure excursion and kept at a constant 10N tension to prevent changes in tendon strain (Figure 1). Retroreflective markers were placed on dissected limbs to track the relative movement of the tibia and tarsometatarsus in 3D across a range of joint angles using a 4-camera Motion Analysis system (300 Hz; Kestrel, Motion Analysis Corporation, Santa Rosa, CA), and automatically synchronized to the linear transducer data within the motion analysis software (Cortex, Motion Analysis Corporation). Joint centres and a mean helical axis were calculated from motion data for each trial and used to calculate flexion angle at the ankle at each timepoint [54]. A cubic spline was fit to the tendon excursion versus flexion angle points using least-squares approximation and tendon excursion was differentiated with respect to angle to estimate moment arm across the measured range of motion (30°-90°). Average values are reported in Table 1.

**Table 1:** Morphological and functional measures by treatment. All values are given as means +/-standard deviations. ‘pVal’ column lists the p-value of statistical comparisons between groups. Bolded rows show statistically significant differences between groups. * indicates data reproduced from Cox et al. (2020).

|  | Restricted | Control | pVal |
| --- | --- | --- | --- |
| Total animals | 8 | 8 |  |
| Body mass kg | 1.7±0.11 | 1.7±0.14 | 0.5 |
| Extensor muscle mass kg | 0.239(0.022) | 0.257(0.02) | 0.18 |
| Average moment arm cm | 0.91±0.05 | 0.94±0.03 | 0.24 |
| Tendon stiffness kN/m | 48.1±13 | 53.5±10 | 0.39 |
| Leg length mm | 345±11 | 349±18 | 0.56 |
| Femur length/leg length | 0.25±0.01 | 0.25±0.01 | 0.18 |
| Tibia length/leg length | 0.36±0.01 | 0.35±0.01 | 0.1 |
| Tmt length/leg length | 0.22±0.01 | 0.22±0.01 | 0.55 |
| Toe length/leg length | 0.17±0.01 | 0.17±0.01 | 0.63 |
| *Jumps/day during growth | 0 | 194(349) |  |
| **At maturity** |  |  |  |
| **\*Max jump takeoff velocity m/s** | **3.3±0.43** | **4.0±0.36** | **0.007** |
| **\*Jump work J/kg** | **37±9.2** | **50±8.4** | **0.013** |
| **\*Peak power W/kg** | **787±165** | **1171±117** | **3.5e-4** |
| **\*Peak jump force/body weight N/N** | **4.7±0.54** | **6.7±0.74** | **5e-5** |

**Figure 1:**
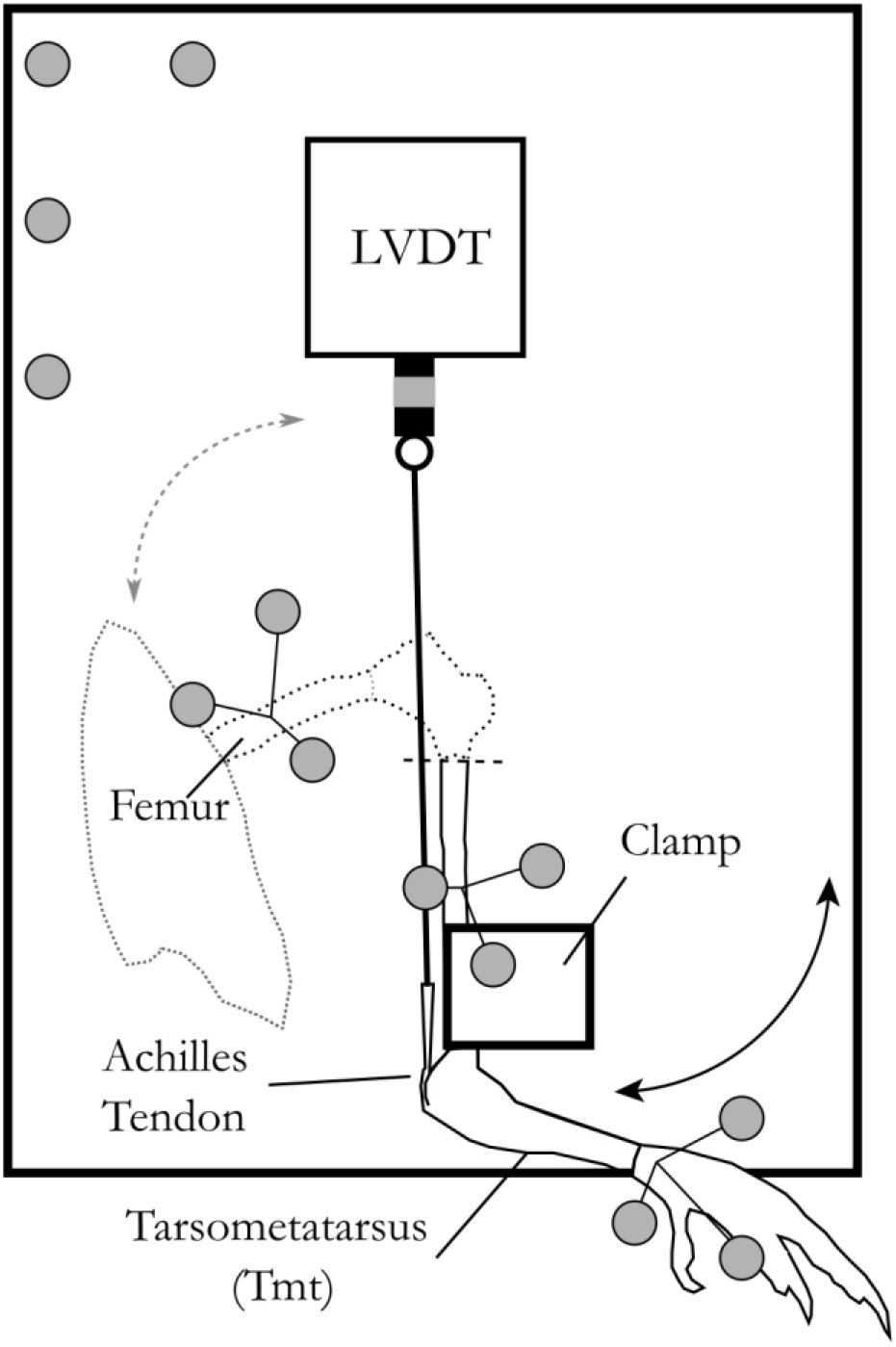
Setup for the tendon travel experiment. The limb was positioned so that the tibiotarsus was held firm by a 3D printed clamp. In the knee joint motion trial, the femur was rotated to move the knee through its ROM. The tibiotarsus was then cut to remove the proximal portion of the limb, allowing for LVDT to be attached to the Achilles tendon. For tendon travel trials, the TMT was rotated to move the ankle through its ROM. Gray coloring represents retroreflective markers on the limb and LVDT. The dotted line outlines the pelvis femur, and knee, which were removed after the knee joint motion trial. The dashed line represents location at which the tibiotarsus was cut after the knee joint motion trial. Figure adapted from Salzano, 2020.

#### Tendon force-length curves

We quantified the tendon force-length properties with material analysis as described in [25]. In short, tendons were detached from the gastrocnemius muscles but left attached at their insertion points on the tarsometatarsus bone. Both the bone and the tendon’s proximal end were connected to a material testing machine (858 Mini Bionix II; MTS Systems Corp; Eden Prairie, MN, United States). Samples were mounted vertically using custom clamps on the tendon aponeurosis and the TMT and attached to a 50-pound load cell (MTS Systems Corp; Eden Prairie, MN, United States). The upper clamp gripped the entire aponeurosis of each sample, leaving only the free tendon exposed to loading. The tendon force-length properties were quantified by loading the tendon cyclically (20 cycles) to 4% strain. The tendon force-length curves were calculated by averaging the data from last 5 cycles of the loading protocol. Tendon force-strain curves were calculated by normalizing displacement by the length of the tendon, *T*_*L*_, measured to the nearest 0.1 mm with calipers while under zero force in the material testing setup. Average values for tendon stiffness given in Table 1 were calculated from the slope of the tendon force-length curve across the last 50 points measured in the last 5 cycles of trials, at strain between 3 and 5%.

#### Tendon Slack Length

The tendon slack lengths for the LG and MG were estimated from experimental measures as described in Appendix A. Because model based estimates of muscle fiber length in a given posture are particularly sensitive to the tendon slack length [55–57] and our calculations involved several simplifying assumptions, we further refined our experimental estimates of tendon slack length by fine-adjusting the tendon slack length parameter in the OpenSim model (see Appendix A for experimental tendon slack length measurement and see below and Appendix B for model development). After experimental moment arms and tendon and muscle properties were added to subject specific models, each model was posed in the individual’s fixed posture. The model’s tendon slack length was adjusted iteratively in the model until the LG and MG normalized fiber lengths were within 1% of the experimentally measured values. These final values are listed in Table 2.

**Table 2:**
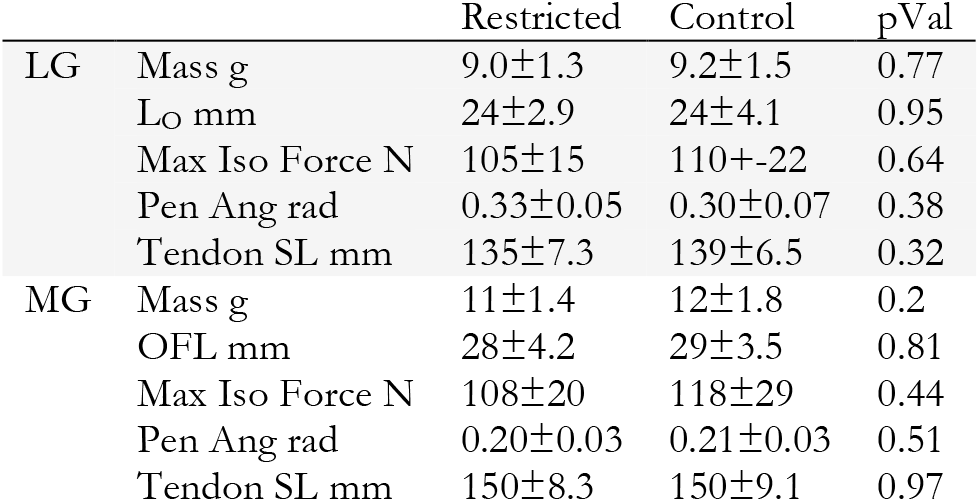
Muscle morphological data by treatment for the lateral (LG) and medial (MG) gastrocnemius muscles: Muscle mass (Mass g), optimal fascicle length, (L_O_ mm), Maximum Isometric Muscle force (Max Iso Force N), muscle pennation angle at optimal fiber length (Penn Ang rad), and tendon slack length (Tendon SL mm). Morphological measures did not vary between groups. All values are given as means +/-standard deviations. pVal column lists the p-value of statistical comparisons between groups.

|  |  | Restricted | Control | pVal |
| --- | --- | --- | --- | --- |
| LG | Mass g | 9.0±1.3 | 9.2±1.5 | 0.77 |
| | $L_O$ mm | 24±2.9 | 24±4.1 | 0.95 |
| | Max Iso Force N | 105±15 | 110±22 | 0.64 |
| | Pen Ang rad | 0.33±0.05 | 0.30±0.07 | 0.38 |
| | Tendon SL mm | 135±7.3 | 139±6.5 | 0.32 |
| MG | Mass g | 11±1.4 | 12±1.8 | 0.2 |
| | OFL mm | 28±4.2 | 29±3.5 | 0.81 |
| | Max Iso Force N | 108±20 | 118±29 | 0.44 |
| | Pen Ang rad | 0.20±0.03 | 0.21±0.03 | 0.51 |
| | Tendon SL mm | 150±8.3 | 150±9.1 | 0.97 |

### Quantifying the influence of restricted jumping on energy storage capacity

We generated a flock of subject-specific musculoskeletal models by modifying the generic model [58] to match experimental values measured for each bird (see Appendix B). With these models, we quantified the capacity of each subject-specific model to store elastic energy in its Achilles tendon across a range of joint postures [Ankle° : 31 to 145, Knee°: −145 to −15, Figure 2 [58])]. At each posture, the simulated LG and MG were activated at 100% and the muscle-tendon unit was equilibrated with the OpenSim MATLAB equilibrateMuscles() function, which adjusts muscle and tendon length such that tendon force and muscle (active and passive) forces balance. The LG and MG insert on the same tendon but, due to OpenSim modeling constraints, these muscles are modeled as having separate tendons. To calculate the stored elastic energy in the combined Achilles tendon, then, we first extracted the resulting force, *f*_*a*_, along the tendon for each muscle. These values were summed and the resulting tendon strain in a single tendon, ∈_*Ta*_, was found from the inverse of the experimentally measured force-strain curve. 

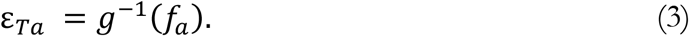

The strain energy stored in the strain of the tendon, *PE* (eq. 10), was calculated by integrating the tendon force-strain curve from zero to the calculated strain and multiplying that by the tendon sample length, *L*_*T*_, as measured at zero strain during the material testing. 

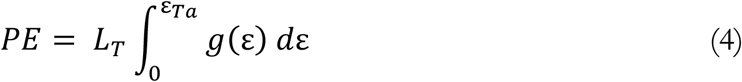

Simulations of 100% activation of the LG and MG were performed across the range of experimentally measured joint angles for the ankle and knee joint (Figures 2A&B). From these simulations, we extracted the maximum elastic energy storage across all postures for each bird and the ankle and knee angles at which the maximum was achieved, and recorded the values in a pre-jump posture (Figure 2C, [31]). Tendon elastic energy stored in the pre-jump posture has been found to be a requirement for the very high power generated in guinea fowl jumping [31,37]. Additionally, we recorded the normalized fiber length for each muscle at this posture at zero activation.

### Statistical Tests

To determine whether components of the gastrocnemius elastic system change systematically in response to changes in demand, we evaluated the influence of treatment group (restricted vs. control) on each element of morphology measured. This was accomplished using t-tests if the homogeneity of variance assumption test was passed and using a Kruskal-Wallis test by ranks when this criterion was not met. Non-parametric analyses are indicated with an asterisk after the p-value in Tables 1&2. The threshold for statistical significance was set at 0.005 after a Bonferroni correction for multiple comparisons. Likewise, the relationship between treatment group and elastic energy storage capacity was evaluated with a t-test after data passed tests for normality and homogeneity of variance, as described above for evaluation of differences between groups of individual elastic system components.

We used stepwise comparison of Akaike information criterion (AIC) values [stepAIC R Mass package [59]] to determine the parameters and coefficients of the full model that best predicted elastic energy storage potential across natural variation of joint postures in preparation for jumps. The full statistical model evaluated included stored strain energy, *PE*, as a dependent factor and, as potential independent variables, tendon stiffness, *tendonK*, the summed maximum isometric force capacity of LG and MG along the tendon, *sumFMax*, the average LG and MG optimal fascicle length, *AvOFL*, and starting muscle length at zero activation of the LG and MG in the pre-jump posture, *AvLenA0c*. We included possible interaction terms between muscle force capacity, tendon stiffness and muscle start length (*sumMaxF*\**tendonK*\**AvLenA0c*) and between optimal fascicle length and tendon stiffness (*avOFL*\**tendonK*) following recommendations by Zajac [60] of functional equivalent muscle tendon joint properties. We did not include muscle moment arm or tendon slack length in the statistical model because they both contributed to the starting muscle length at any given joint posture.

To quantify the relative explanatory power of morphological variation of any individual element to predict stored energy to a systems level approach, we compared individual parameter models to the best multi-parameter model found by stepwise comparisons described above. The AIC value of the best model was compared to AIC values of models with individual predictors and their relative explanatory power computed [61].

The relationship between Achilles tendon elastic energy storage capacity and experimentally measured muscle-mass-normalized peak power output and jump work were both tested with a linear model with elastic energy storage as the dependent variable and peak power or jump work as the independent variable. See [62] for details on how power and work were calculated from force plate data.

## Results

### Variation in individual elements of the elastic system

We found no statistically significant differences in any individual morphological property between birds that jumped during growth and those that did not (all p>= 0.1, Tables 1&2).

### Energy storage capacity between groups

We found no significant differences in the capacity to store energy in the strain of elastic elements between birds restricted and unrestricted from high power activities during maturation, despite small differences in tendon strain that did not reach significance (Table 3, Figure 3). This held true both at the peak crouched posture before jump initiation (Figure 2C, p=0.43) and at the posture that optimized elastic energy storage (Figure 2D, pVal=0.44). It should be noted that the optimum posture for elastic energy storage was at the most extended knee angle and the most flexed ankle angle tested and was ∼85° more extended ankle angle than birds used in preparation for a jump. While this more extended posture lengthened the gastrocnemius and increased the energy stored in the Achilles, it shortened the operating length of the knee extensor muscles, reducing their force-generating capacity. In the prejump posture, the shorter gastrocnemius length decreases elastic storage capacity by 12% for control birds and 10% for restricted birds (Table 3) in comparison to the optimal posture for energy storage.

**Table 3:** Comparison of energy storage capacity and normalized fiber length between restricted and unrestricted birds at maturity across all postures and in the pre-jump posture (shaded grey). Normalized muscle length at the start and end of contraction designated by Average n.Fiber length at A0 and A100 respectively.

|  |  | Restricted | Control | pVal |
| --- | --- | --- | --- | --- |
| At max strain<br>energy posture | Energy Storage Potential J | 0.3±0.065 | 0.26±0.12 | 0.44 |
|  | Tendon Strain | 0.12±0.016 | 0.098±0.021 | 0.031 |
|  | Average n.Fiber length at A0 | 1.1±0.051 | 0.99±0.098 | 0.11 |
|  | Average n.Fiber length at A100 | 0.75±0.051 | 0.72±0.054 | 0.27 |
|  | Knee Angle at Max PE° | -131±1.9 | -130±0 | 1 |
|  | Ankle Angle at Max PE° | -45±0 | -45±0 | 1 |
| In pre-jump<br>posture (Knee:<br>-135 Ankle:<br>120) | Energy Storage Potential J | 0.27±0.055 | 0.23±0.11 | 0.43 |
|  | Tendon Strain | 0.12±0.015 | 0.094±0.021 | 0.028 |
|  | Average n.Fiber length at A0 | 1±0.05 | 0.96±0.097 | 0.11 |
|  | Average n.Fiber length at A100 | 0.73±0.048 | 0.7±0.051 | 0.28 |

**Figure 3:**
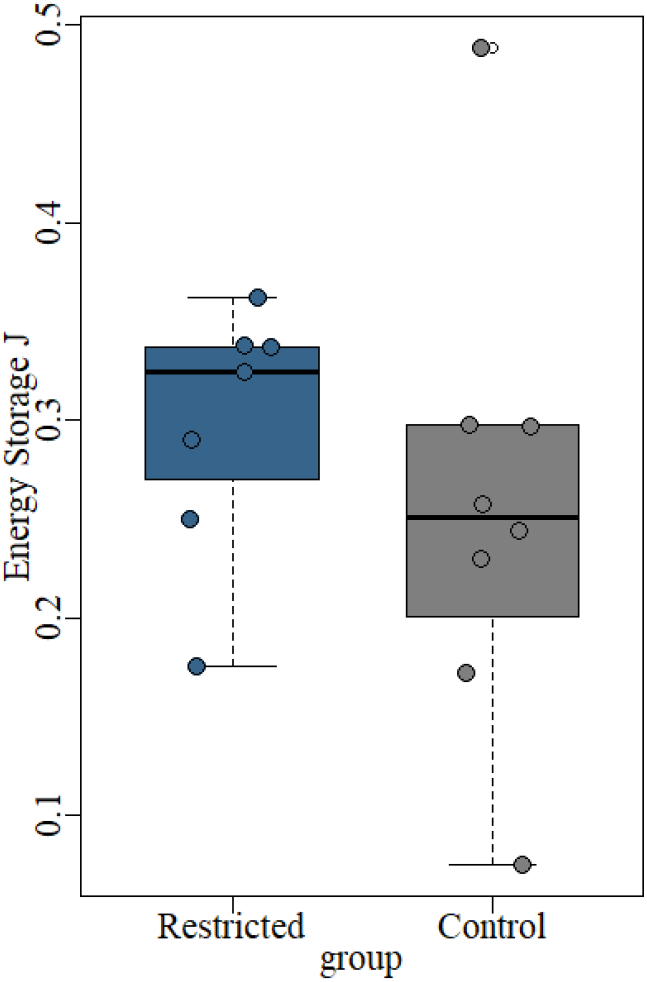
Restricting high power activities does not decrease or significantly influence the capacity to store elastic strain energy. Each dot represents data from one individual.

### Morphological predictors of elastic energy storage capacity

We found that the amount of energy stored in strain of the tendon was best explained by variation in the average of the passive LG and MG muscle length at activation onset, *avLenA0* (Table 4). Figure 4A illustrates that tendon strain energy increases at longer starting lengths both between individuals and, even more strikingly within individuals, across postures. Muscle force capacity along the tendon was the next most explanatory variable and, like normalized muscle length, shows a positive relationship with energy storage (Figure 4B). In contrast, longer muscle optimal fascicle lengths reduced energy storage (pVal: 0.03, Table 4, Figure 4D) when evaluated as an induvial predictor, but was not a significant factor as a predictor in the full multi-parameter model. Opposite to our predictions, tendon stiffness did not significantly correlate with elastic strain energy when evaluated as an individual predictor (pVal>0.1 Table 4, Figure 4C), but did improve the explanatory power of a full model. The stepwise AIC comparison of full and reduced models found the sum of LG and MG maximum force capacity along the tendon, *sumFMax*, average optimal fascicle length, *avOFL*, muscle length at activation onset, *avLenA0*, tendon stiffness and the interaction of tendon stiffness and muscle force capacity as the independent predictors that best correlated with stored elastic strain energy. The relative explanatory power of each predictor following similar patterns as seen in the individual analyses with muscle start length and force capacity showing the greatest predictive power. This full model had an R^2^ of 0.93 and was over 280,000 times more likely to explain the variation in strain energy than any model with only one explanatory variable.

**Table 4:**
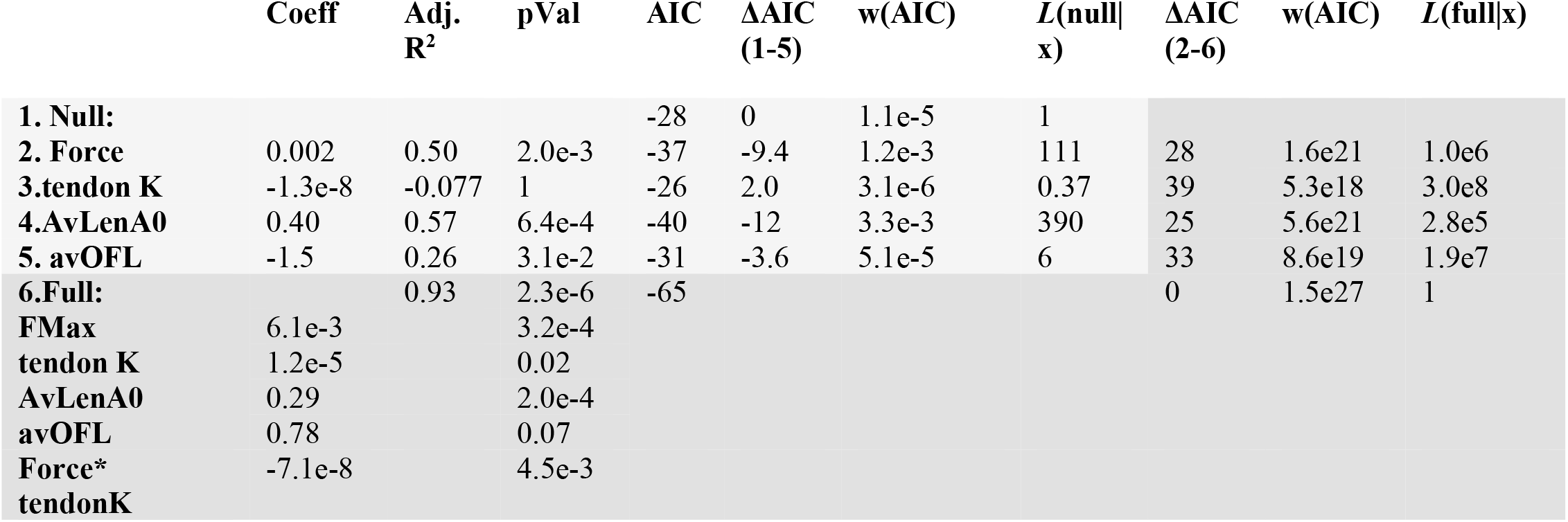
Results of models comparing how well variation of individual elastic elements explain variation in elastic storage potential (dark shaded grey regions) and stepwise model comparisons of models with multiple predictors (lighter shaded region) find that muscle operating length at the start of activation (AvLenA0) is most predictive of energy storage, with muscle force capacity, the sum of lateral and medial maximum isometric force (Force), and average gastrocnemius optimal fiber length, avOFL, also significant.. Tendon stiffness, Tendon K, add little to no additional predictive information. A multi-predictor model (Full:, darker shaded region), explained variation in energy storage capacity over 160,000 times better than any individual predictor. Likelihood comparisons between the null and individual models are designated by L(null|x), and between the full model and individual predictor models by L(full|x). Akaike weights are listed under w(AIC)

**Figure 4:**
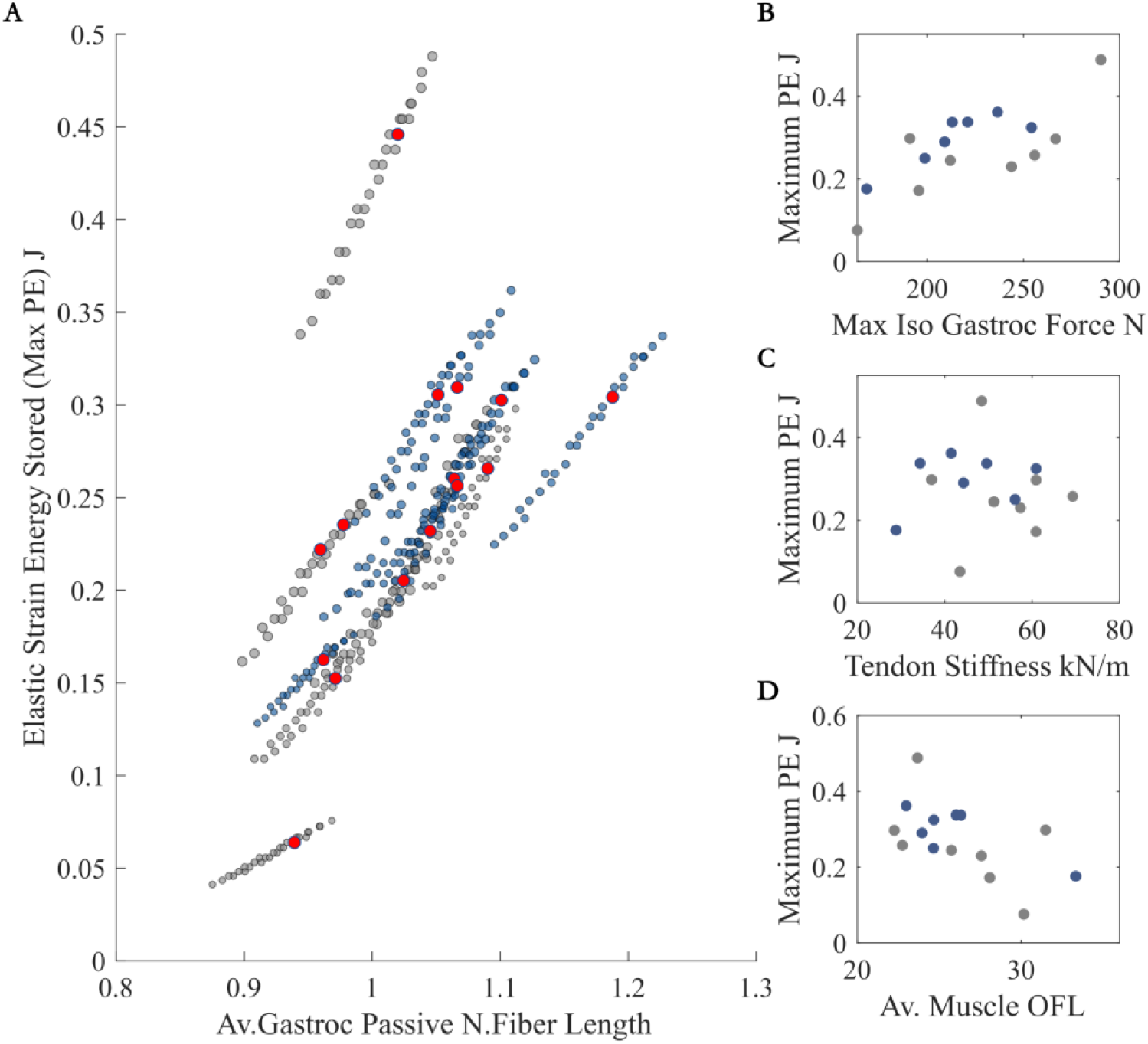
Energy stored in the tendon increases with passive normalized fiber length (length at onset of muscle activation) across different postures (A) and muscle force capacity (B), decreases with muscle optimal fiber length (D), but did not consistently vary with tendon stiffness (C). The variable that most predicts elastic strain energy is muscle normalized fiber length at activation onset (A), with muscles operating at longer lengths enabling greater elastic energy storage. Color designates group (blue: restricted, grey: control). In A), red dots identify the strain energy stored and normalized fiber length in the pre-jump posture. In B, C, and D each marker represents data from one individual.

### Energy storage capacity vs jump performance

We found little to no correlation between energy storage capacity predicted by simulations and experimentally collected jump metrics of either muscle mass-specific work or power (Figure 5). A linear model showed no significant relationship between either peak power jump power per kg of muscle mass capacity (t = −036, p = 0.72, Adj R^2^=-0.07) or jump work (t = −0.05, p = 0.96, Adj R^2^=-0.08) and strain energy in the pre-jump posture. The negative adjusted R^2^ values for both tests shows that the variation in jump work or peak power explains only a negligible amount of variation in elastic energy storage potential. The scatterplot of standardized predicted values versus standardized residuals for both variables showed that the data met the assumptions of homogeneity of variance and linearity and the residuals were approximately normally distributed.

**Figure 5:**
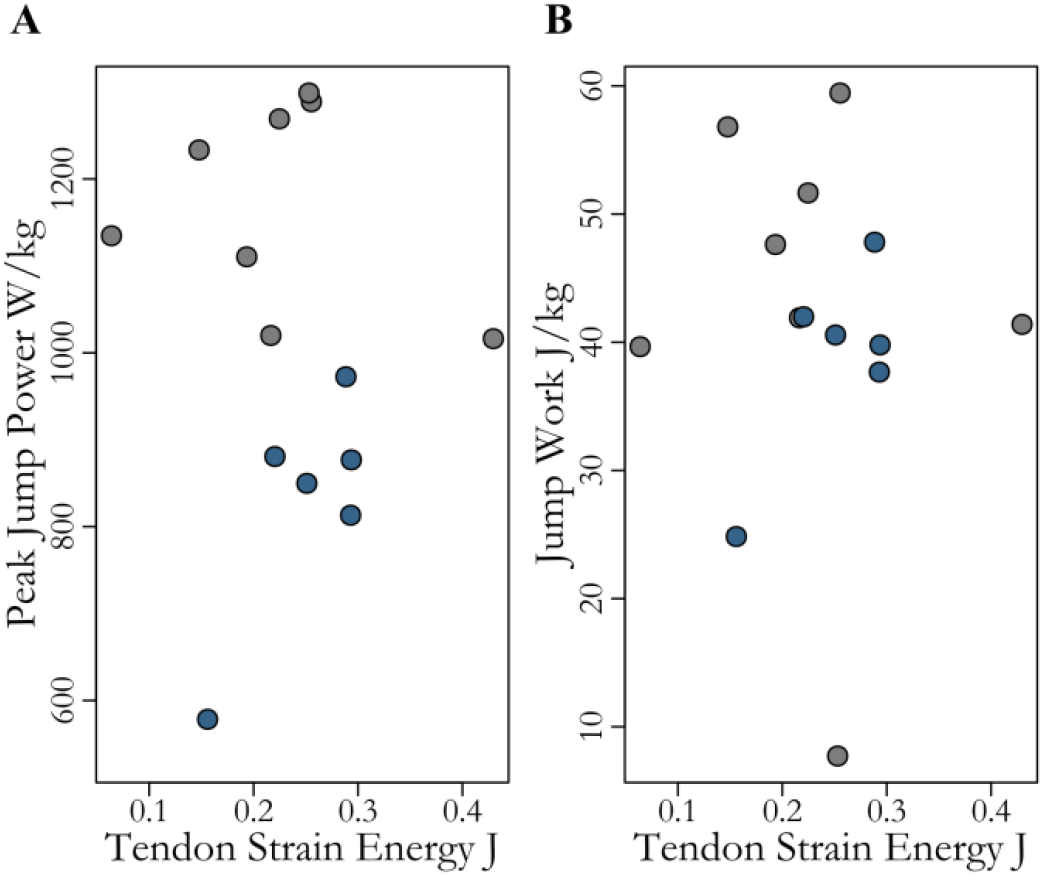
Neither peak jump power nor jump work increase systematically with maximum tendon strain energy across all individuals. Markers designate data from one individual in pre-jump posture and color differentiates treatment groups (grey: control, blue: restricted)

## Discussion

We found the gastrocnemius elastic system of the guinea fowl robust to variations in locomotor conditions during growth. Neither properties of individual components nor energy storage capacity varied between groups of birds which did and did not jump throughout maturation. Nor did we find any correlation between energy storage capacity and jump performance. Variation in muscle operating length across individuals predicted energy storage capacity better than any fixed morphological property and a systems approach incorporating multiple components was substantially able to predict energy storage capacity better than variation along any individual element.

### Do components of the gastrocnemius elastic system change systematically in response to changes in power and work demand during growth?

Contrary to our predictions, we saw, in general, no systematic changes between the gastrocnemius elastic system in response to decreased demand for high power and work activities during maturation. Surprisingly, birds that were restricted from jumping throughout their entire growth period (Table 1) developed elastic systems that were largely indistinguishable from the control group that jumped, on average, almost 200 times a day and exhibited greater peak jumping power and work at adulthood (Tables 1&2). Nor did we see differences in body mass, extensor muscle mass, tendon stiffness or leverage (moment arm and/or bone lengths).

Two factors may account for why we saw no systematic changes in the elastic system while observations of morphological plasticity in response to changes in functional demand abound [63– 67], even in guinea fowl in particular [43,68]. First, several studies suggest that plasticity may vary by life stage, with lower or inconsistent plasticity in growing animals [69–71]). Thus, the inconsistency between the lack of plasticity in our study and the morphological variation found by others suggests that guinea fowl may exhibit lower plasticity during maturation than in adulthood. Fast-growing species, like guinea fowl, might outpace environmental fluctuations with rapid growth and not invest in developmental plasticity [72,73]. A slow growing species (humans for example) might have a selective advantage with greater developmental plasticity. Thus, while our treatment may have been powerful enough to induce morphological changes in adults, rapidly growing guinea fowl may be more robust to environmental perturbations.

Second, our results could be consistent with the results of other studies if the plastic response to decreased demand is not inferable from changes in response to increases in demand. For example, it may be that the increase in muscle mass that occurs in response to a certain *increase* in functional demand is greater than the decrease in muscle mass that occurs in response to the equivalent *decrease* in demand. Many studies find clear evidence of morphological plasticity, but this was in response to increased mechanical load [24,43,66,68,74,75] and extreme disuse [76–78]. Our intervention, however, eliminated jumping and while maintaining consistent low intensity exercise (i.e. walking) and thus we did not induce chronic offloading, as had been the goal in several previous disuse studies. Our results suggest that there may not be a linear dose-response relationship between changes in functional demand and morphological variation. Instead, as recently suggested [25], there may be a range of variation in demand that is not extreme enough to induce physiological or morphological modification above those under developmental control. If this region of stasis encompasses a wider range of disuse, it could explain both why offloading studies often require extreme disuse, like bedrest or limb immobilization [76,77,79], to induce change and why we found no systematic morphological changes here. Eliminating jumping may not be an extreme enough disuse signal to induce musculoskeletal plasticity.

Thus, while we found no systematic morphological variation when restricting high power activities during maturation, this does not necessarily imply that the morphology of elastic systems does not plastically adapt to variations in functional demand. But it does suggest that there are conditions in which elastic systems may be insensitive to functional variation.

### Is the energy storage capacity reduced in individuals that did not jump during growth?

Despite this lack of consistent morphological variation between our treatment groups, restricted birds generated lower absolute and muscle-mass-specific power and work during jumping. This suggests either that small morphological changes in individual elastic elements compound to alter elastic system function, that variations are significant in other MTU’s that we did not quantify, or that behavioral or neural variation account for the difference in jumping performance. Our systems level analysis aimed to specifically address the question of whether morphological variation compound within the elastic system to enable unrestricted birds to store more energy in their Achilles tendon in preparation for a jump. Again, contrary to our predictions, simulations in our subject-specific models resulted in no differences between groups in their ability to store elastic energy. Taken together, the finding of minimal changes to individual muscle-tendon unit components, and no effect on the overall elastic energy storage, could indicate that morphology necessary to enable jumping is highly conserved. This could happen if rapid movements are very critical to fitness, as may be the case for prey animals for whom evasion is critical.

### Which type of morphological variation has the greatest influence on energy storage capacity?

The first two analyses focus on plasticity of elastic systems and quantified the influence of rearing conditions on the morphology of individual components and how that variation influenced energy storage capacity across treatment groups. Our last two analyses utilize the variation within and across our treatment groups to further probe the relationship between form and function in elastic systems.

Evaluating the best predictors of energy storage, we found that muscle properties far outweighed the influence of tendon stiffness. Surprisingly, maximum isometric muscle force, while correlating with energy storage, was not the most important factor. Instead, normalized muscle length at the start of contraction was the best individual predictor of energy storage, with the longest normalized muscle lengths enabling greatest elastic storage [(in agreement with results from [28,80]]. This may be because muscles that start contracting on the descending or plateau region of the force length curve increase force capacity as they shorten, resulting in a greater equilibrium force, while muscles starting on the ascending limb of the force-length curve lose force capacity as they shorten against a tendon during an isometric contraction [58]. Further, we were particularly surprised to find that tendon stiffness alone had little to no predictive power of energy storage. Together these data suggest that, between individuals or across an individual’s lifetime, the large variation in force capacity due to force-length or force velocity effects may overshadow the influence of variation in tendon properties in determining tendon strain energy. This conclusion is consistent with studies in humans that found no correlation between tendon stiffness and vertical jump height [40]. Yet, this idea runs contrary to the focus on spring properties [23,27,39,81–85] or relative spring and maximal muscle properties [3,75] in many studies that try to connect form and function in elastic systems. Our results suggest that instead, between or within individuals, elastic energy storage capacity may be more sensitive to variations that alter muscle operating lengths (tendon slack length, optimal fascicle lengths, joint postures) or cross-sectional area than changes in the tendons themselves.

Further, our results also highlight the importance of analyzing the components of an elastic system in concert rather than trying to infer performance from variation in one component. Our full model that included both muscle (max muscle force and starting length) and tendon properties explained changes in energy storage capacity over 280,000 times better than variation in any individual property, even when penalizing models for complexity. This, again, emphasizes the limitations of reductionist approaches to understanding how musculoskeletal morphological variation influences the energy storage capacity of an elastic system.

### Does elastic energy storage capacity predict peak jump powers and work?

The finding that maximum energy storage capacity of the gastrocnemius elastic system did not predict jump performance further supports the conclusion of the need to switch our focus from individual elements to analyzing the elastic system encompassing both morphology and neural control. Contrary to our expectations, individuals who developed elastic systems capable of storing greater energy in their tendons did not take advantage of that ability to produce more powerful jumps. This suggests that morphology may play a smaller role than neural control in determining contribution of elastic energy storage in jumping. Musculoskeletal morphological variation may not be the main factor limiting jump performance.

The interaction between tendon and muscle force-length curves may, in part, provide a mechanistic explanation for this disconnect between morphology and performance. Here we include a conceptual diagram to illustrate this (Figure 6). If we plot the muscle and tendon force-length curves on the same figure, it is possible to visualize how they might interact. If the muscle operates on the ascending or plateau region of the force length curve where passive forces are minimal, and we assume that there is no slack in the tendon, a tendon strain of zero will coincide with the muscle length at the start of a contraction. Any tendon strain, then, is equal and opposite to the change in muscle length. During a fixed end contraction, the maximal tendon strain occurs when the tendon force equals the total muscle force of the three heads of the gastrocnemius (empty circles, Figure 6). The tendon strain at equilibrium, then, is dramatically influenced by the length of the muscle when it begins to contract (Figure 6A), reaching higher values for contractions starting at longer muscle lengths. As our results suggest (and as can be visualized by comparing the differences in the areas of the grey shaded regions between Figure 6A and B), these dynamics can be larger than the influence of naturally occurring variations in maximal force capacity (Figure 6B). And while muscle operating lengths are constrained by morphology (OFL, pennation angle and moment arms), they are also easily varied with joint angle. Large variations in maximum muscle force could be compensated for by small variations in posture largely under neural control. Thus, as our results suggest, dynamic factors like muscle operating length may influence energy storage more than temporarily fixed musculoskeletal features like maximum muscle force capacity.

**Figure 6:**
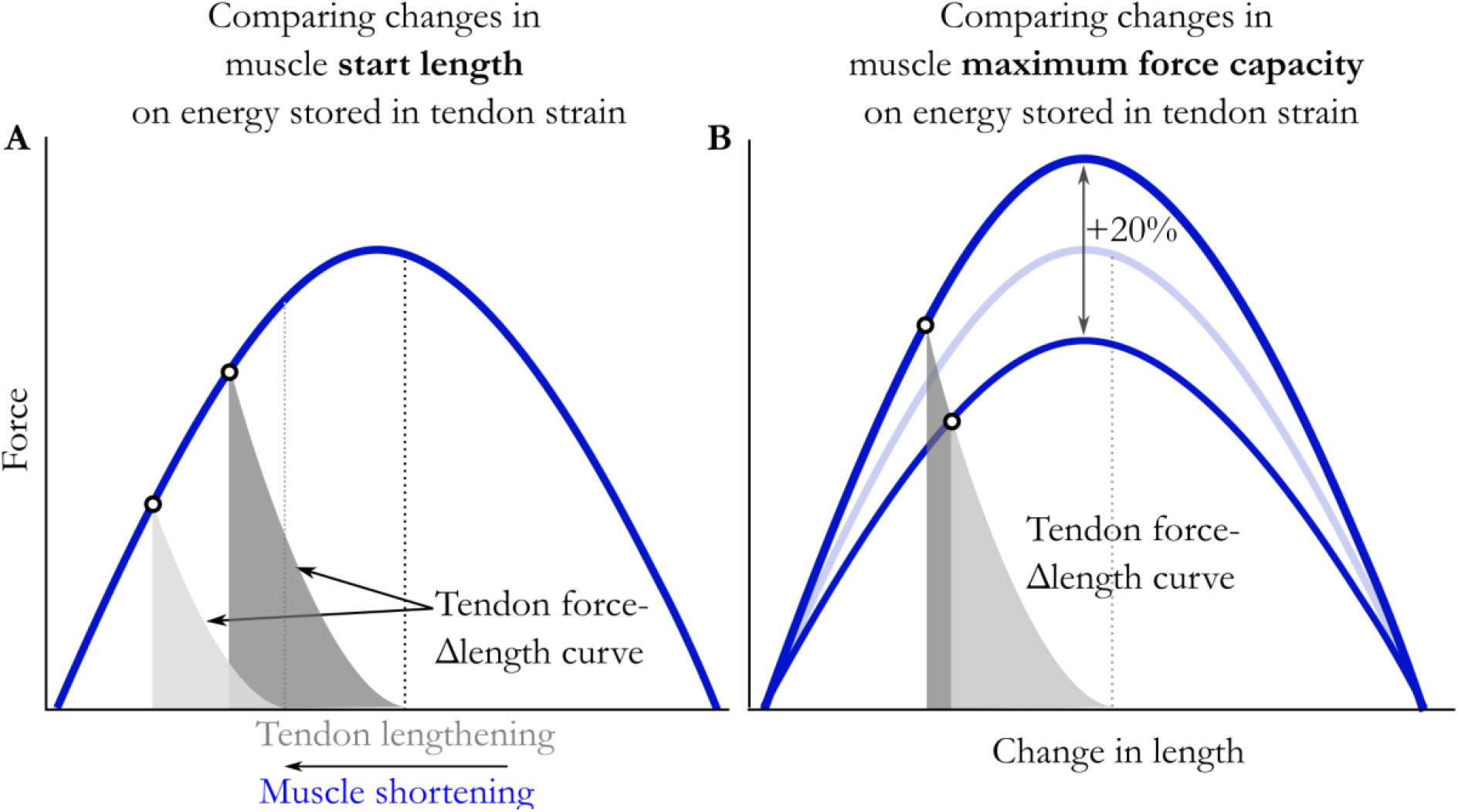
The energy stored in the tendon during a fixed end contraction is constrained by the interaction of the muscle (blue curve) and tendon force-Δ length curves (grey curve). Since tendon length changes must equal and opposite to muscle length changes, we can plot them at the same scale. Changes in muscle length at the onset of contraction (A) can have a larger influence on energy storage than variations in muscle force capacity (B). The variation in maximum muscle force depicted here represent variation observe in our subjects. Here changes in start length (‘0.9-1.2) and muscle force capacity (±10%) reflect experimentally measured variation in our population.

This suggests that there may be a large range of morphological variation that can be compensated for with neural plasticity and brings up questions of how the two interact. Does variation along a particular morphological axis correlate with systematic changes in neural control? If so, how do individuals search though the neural possibility space? What are the limits of neural compensation? Do we see greater morphological plasticity of the components of elastic systems in conditions that push the limits of neural plasticity?

### Potential interactions between musculoskeletal and neural variation in elastic energy storage

This possible sensitivity of performance to muscle operating lengths also suggests directions for further studies exploring the interaction between morphological and neural or kinematic variation. Here we assumed a uniform pre-jump posture across individuals based on measured average values [31]. Yet our results suggest variations in posture between individuals may play an important role in understanding the relationship between form and function in elastic systems. For instance, one could explore whether individuals with shorter muscle operating lengths adopt a less flexed posture to enable greater tendon energy storage in preparation for a jump. Likewise, the addition of a counter movement proceeding a jump, not common in guinea fowl [31,37] but present in many other species [86,87], could minimize influence of muscle constraints on energy storage. This dependence of performance on the interaction between morphology and neural control again reinforces the need to expand our scope from that of the muscle-tendon unit to that of the elastic system.

Therefore, while musculoskeletal morphology may set the bounds of possible energy storage, individuals may not operate at their limits of elastic potential. This suggests either a significant behavioral component (restricted birds simply may have not tried as hard to jump) or that there may be benefits to real-time tunability in elastic systems. The jump of a guinea fowl is powered both by tendon recoil and simultaneous muscle work [31], as is common in many larger animals [86–89]. The muscles that load the tendon pre-jump also contract during takeoff to contribute power to the jump. In these hybrid systems, trade-offs between maximizing tendon strain energy and muscle power may explain our findings that the pre-jump posture of guinea fowl did not optimize energy storage in the tendon. Adjusting muscle lengths to maximize tendon strain may hamper muscles fiber work during takeoff. Further, in a complex system such as this, with dozens of individual muscle-tendon units spanning multiple joints and working in concert with direct drive muscles with little tendon, the difference between a great jumper and a good jumper might depend less on the maximal storage capacity of any one muscle tendon unit (i.e. its musculoskeletal morphology) and more on fine adjustments of neural control to harmonize the output of the collective system [90–92].

Together, our experimental and modeling analyses suggest that performance advantage of the control birds, who practiced jumping throughout maturation, may lie less in the body’s modification of individual elastic elements, and instead, in the fine tuning of neural circuits to coordinate muscle activation timing to take better advantage of what they each possess. While restricting normal locomotor behavior during growth (i.e., eliminating practice) likely leads to deficits in neural control, neural plasticity is potentially a rapidly reversible pathway to adapt an elastic system to functional variation. Given the potential short timescale of neural plasticity [92,93], greater sensitivity of neural locomotor/movement stimuli could allow the individual to adjust the dynamics of an elastic system during growth without making potentially irreversible changes to morphology that could be detrimental in subsequent stages of growth or in adulthood if environmental conditions or functional demand rapidly change. Thus, one could interpret the results of our study as suggesting that practice during growth may indeed be more related to forming the neural framework for jumping than for forming the musculoskeletal framework. This also suggests the specific hypothesis that individuals restricted from an activity during growth may be capable of reversing the resulting neural deficits with practice later in life.

### Limitations

Several modeling simplifications could have influenced our results. For instance, the gastrocnemius elastic system is not the only one that could contribute to jump power. While the Achilles is the largest tendon involved, many other digital tendons spanning both the ankle and tarsometatarsus joint have the potential to contribute to jump power but were not included in our analysis Furthermore, changes in the characteristics of muscles spanning proximal joints may also have contributed to the differences in jump power but these muscles were not modeled. Additionally, potentially important dynamic effects were ignored. For simplicity, we simulated the amount of energy stored in the Achilles tendon during fixed-end contractions, where the joint posture was constant as the muscle and tendon dynamically responded to increasing muscle activation.

Activating muscles while altering joint posture would alter these dynamics, perhaps amplifying the influence of individual differences in input or output lever lengths or force-velocity effects not apparent from group averages. Likewise, we did not measure and include individual variation in muscle/aponeurosis passive elastic properties that could significantly alter energy storage [36,94,95]. While this is a common approach in musculoskeletal modeling [50,60,96], variation in the aponeurosis and free tendon stiffness [97,98] have the potential to introduce errors [16,99].

Another possible limitation was the modeling choice to focus on the potential for an individual to store energy in the strain of their tendon. How that energy is released and how that energy release interacts with synchronous muscle activation could also influence jump performance [5]. Because jumps are likely powered both by tendon recoil and muscle work [31,80], there may be a tradeoff between the work the muscle puts into tendon strain and that which is left available to power the jump during tendon recoil [100]. Future work could involve simulation of jumps in these subject-specific models to assess the contributions of these dynamic factors. Thus, while we found no consistent change in components of the gastrocnemius elastic system due to decreased demand for high power activities during growth, more complex models may provide insight into the ways in which morphological variation constrains performance.

### Summary

We found that decreasing the demand for high power and work during growth can influence adult performance but does not necessarily lead to morphological plasticity. We found no difference in energy storage capacity between groups which did and did not jump throughout maturation or any correlation with experimentally measured jump performance. We conclude that gastrocnemius elastic system in the guinea fowl displays little to no morphological plastic response to decreased demand during growth and that neural control of elastic systems may constrain performance more than morphology.

## Author Contributions

SMC, JR, SJP, MQS contributed to the conception and design of the study. JR, MQS, SJP and SMC developed methodologies and KK, AD, MQS, SMC collected the data. SMC, MQS analyzed the data and SMC prepared the figures. SMC and JR drafted the initial manuscript. All authors contributed to manuscript editing and approved the final manuscript.

## Funding

This study was supported in part through a seed grant from the Center for Human Evolution and Diversity, The Pennsylvania State University, and through the National Institute of Arthritis and Musculoskeletal and Skin Diseases of the National Institutes of Health under grant number R21AR071588. The content is solely the responsibility of the authors of this paper and does not necessarily represent the views of the National Institutes of Health.

## Conflict of Interest

The authors declare that the research was conducted in the absence of any commercial or financial relationships that could be construed as a potential conflict of interest.

## Data Availability

TBD

## Appendix A

### Estimation of tendon slack length from experimental measures

Tendon slack length was estimated from experimental measures of muscle and tendon morphology as follows. First, the maximum isometric force along the tendon was calculated from the maximum force along the fiber and the pennation angle at optimal fascicle length, *θ*_*OFL*_, according to the equation: 

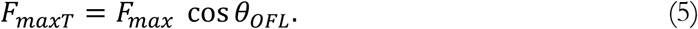

The passive force of the muscle exerted on the tendon in the experimentally measured posture was found from the normalized passive muscle force as a function of normalized fiber length curve, *f*_*np*_(*nFl*) [101,102]. This was first scaled, for each muscle, by the maximum isometric force along the tendon, *F*_*maxT*_, 

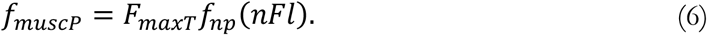

By normalizing the experimentally measured average fiber length by the muscle’s optimal fascicle length, we could calculate the normalized fiber length of the muscle in the fixed posture, *nFl*, allowing us to solve equation (4) for the passive force, *f*_*muscP*_, each muscle exerted on the tendon in the experimental posture. Since the three heads of the gastrocnemius attach to the Achilles tendon, the passive force of each muscle was calculated separately and summed. As the gastrocnemius intermedia head makes up ∼10% of the total gastrocnemius muscle by volume, the passive contribution of this muscle was not experimentally determined for each bird but was estimated from values previously collected [58]. The passive force exerted by the muscle must be balanced by an equal tendon force, thus, the summed passive muscle forces equal the passive force the tendon experienced in the experimental posture.

The MTU lengths, *L*_*MTU*_, were measured on the fixed limbs by digitizing the three-dimensional paths of the MG and LG from their origins on the tibiotarsus and femur, respectively, to the insertion of the Achilles tendon on the hypotarsus. This approach inherently includes the aponeurosis in the overall tendon length. Digitizing was done using a digitizing arm (Microscribe 3DX, Immersion, San Jose, CA). The MTU path was described by 11 points. The linear distances along the MTU path were summed to obtain an overall MTU length. This experimentally measured MTU length, *L*_*MTU*_, is the sum of the measured fiber length, *L*_*M*_, the tendon’s slack length, *L*_*T*_, and length change in the tendon (tendon stretch) due to passive muscle fiber force. The length change in the tendon due to passive muscle fiber force can be described as the tendon strain, ε_*T*_, times its tendon slack length, *L*_*T0*_. 

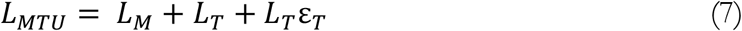

The strain in the tendon due to the passive muscle fiber force, ε_*T*_, was calculated using the experimentally measured tendon force-displacement curve. The tendon force-displacement curve was normalized by tendon length to generate a force-strain curve. 

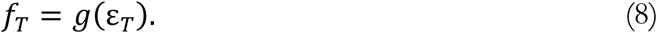

The strain at which the tendon force is equal to the passive fiber force can then be found from the inverse of equation (6) and the passive muscle force, *f*_*MP*_. 

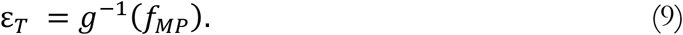

The tendon slack length, for each muscle, then, can be calculated from equations (5) and (7). 

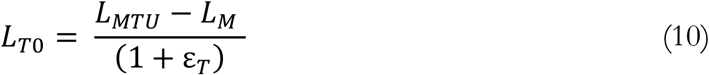

## Appendix B

### Development of subject specific models

To perform system level analyses, we modified the generic OpenSim guinea fowl model [58] to generate subject-specific models for each individual. First, the generic model was scaled to match the measured bone lengths and body mass for each bird and saved as distinct models. In each subject specific model, the generic LG and MG maximum isometric force, pennation angle, optimal fascicle length and tendon slack length were modified to match the experimentally measured and calculated properties.

The moment arms of the LG and MG acting at the ankle was fit to experimental values by adjusting the of the size and orientation of the cylindrical wrapping surface for the Achilles at the ankle. During the trial-and-error fitting process, the radius, translation, and rotation of the wrap surface was modified, and the resulting moment arm was compared to the experimentally collected data at 31-34 points across the experimental range with a mean moment arm normalized root mean square of the error (Figure S1A) of 0.009±0.007.

**Figure S1:**
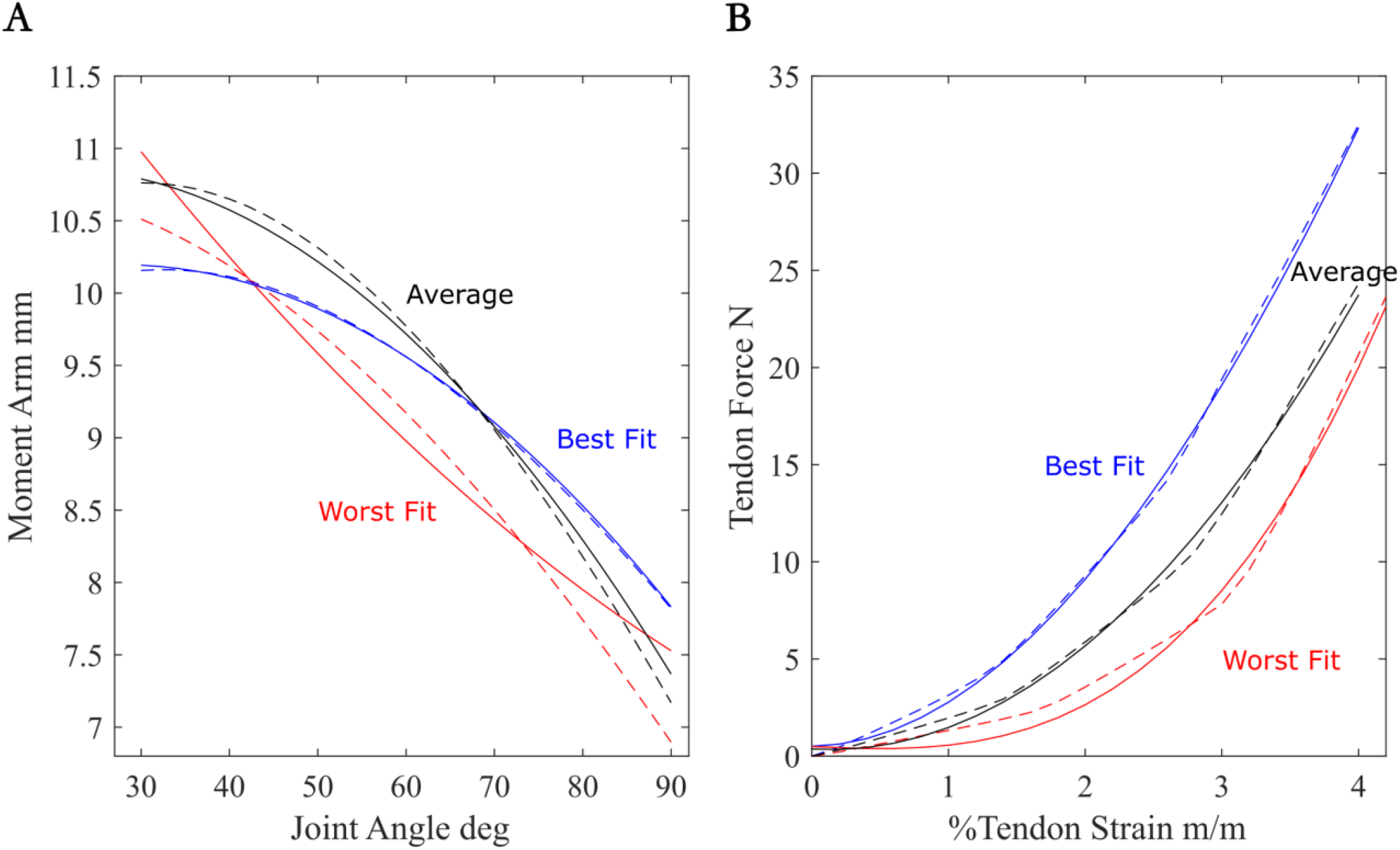
Example comparisons between experimental (solid lines) and modeled (dashed lines) moment arms (A) and tendon force-strain curves (B). In each plot, experimental and modeled curves are displayed for three animals, showing the best, the average and the worst fit across individuals.

Additionally, the tendon force-strain curve was updated to match experimentally collected force-strain values. Because OpenSim scales the tendon force-strain curve by the maximum isometric force of the muscle, each tendon force-strain curve was normalized by the maximum isometric force capacity of the LG or MG, respectively. The parameters of the Millard muscle model’s tendon force-strain curve were iteratively varied for both the LG and MG and compared to the experimental curve for each individual, resulting in an average root mean square error (normalized by tendon force) over 26-31 points for each tendon force-length curve (Figure S1B) of 0.061+0.025.

## Notes

### Competing Interest Statement

The authors have declared no competing interest.

